# Gene expression profiling reveals U1 snRNA regulates cancer gene expression

**DOI:** 10.1101/099929

**Authors:** Zhi Cheng, Yu Sun, Xiaoran Niu, YingChun Shang, Zhenfeng Wu, Jinsong Shi, Shan Gao, Tao Zhang

**Author notes:** These authors contributed equally to this paper (Zhi Cheng, Yu Sun). The corresponding authors. SG, TZ.

## Abstract

U1 small nuclear RNA (U1 snRNA), as one of the most abundant noncoding RNA in eukaryotic cells plays an important role in splicing of pre-mRNAs. Compared to other studies which have focused on the primary function of U1 snRNA and the neurodegenerative diseases caused by the abnormalities of U1 snRNA, this study is to investigate how the U1 snRNA over-expression affects the expression of genes on a genome-wide scale. In this study, we built a model of U1 snRNA over-expression in a rat cell line. By comparing the gene expression profiles of U1 snRNA over-expressed cells with those of their controls using the microarray experiments, 916 genes or loci were identified significantly differentially expressed. These 595 up-regulated genes and 321 down-regulated genes were further analyzed using the annotations from the GO terms and the KEGG database. As a result, three of 12 enriched pathways are well-known cancer pathways, while nine of them were associated to cancers in previous studies. The further analysis of 73 genes involved in 12 pathways suggests that U1 snRNA regulates cancer gene expression. The microarray data with ID GSE84304 is available in the NCBI GEO database.

## Introduction

U1 small nuclear RNA (U1 snRNA) is one of the most abundant noncoding RNA [1]. U1 snRNA has a length of 164 nt in human and its protein binding sites are highly conserved in insects and mammals [2]. The primary function of U1 snRNA is its involvement in the splicing of pre-mRNAs in nuclei. It has been known that snRNAs are synthesized in nuclei and transported to cytoplasm to retrieve other core spliceosome components, then return to nuclei for their functions [3]. U1 snRNA also functions in the regulation of transcription factors or RNA polymerase II, and maintaining telomeres.

The abnormalities of U1 snRNA can cause defects in pre-mRNA splicing, which are considered as a primary cause of human diseases [4]. In the year of 2013, Bai *et al.* discovered the cytoplasmic aggregation of U1 snRNA with several U1 small nuclear ribonucleoproteins (U1 snRNPs) in Alzheimer’s Disease (AD), but not in other examined neurodegenerative diseases including Parkinson's Disease (PD) and Frontotemporal Lobar Degeneration (FTLD) [5]. Bai *et al.* also demonstrated that the cytoplasmic aggregation of U1 snRNA and U1 snRNPs resulted in a loss of nuclear spliceosome activity, which altered the expression of the Amyloid Precursor Protein (APP) and the Amyloid Beta (Aβ) protein. APP includes a region that generates the Aβ protein, the aggregation of which has been considered as a main factor to cause AD. In the past 20 years, the studies of U1 snRNA have focused on its primary function, particularly related to neurodegenerative diseases caused by the abnormalities of U1 snRNA. Besides the primary function, a non-canonical role for U1 snRNPs has been reported to protect pre-mRNAs from drastic premature termination by cleavage and polyadenylation (PCPA) at cryptic polyadenylation signals (PASs) in introns [6] [7]. Although the previous studies demonstrated this new function by knockdowning U1 snRNA, no results of U1 snRNA over-expression had been yet reported in published papers before this manuscript was submitted.

In this study, we built a model of U1 snRNA over-expression in a rat cell line. Based on this model, we compared the gene expression profiles of U1 snRNA over-expressed cells with those of their controls to reach two research goals: 1) to study how the U1 snRNA over-expression affects the expression of genes on a genome-wide scale and identify the significantly differentially expressed (DE) genes; 2) to globally investigate the cellular effects brought by these DE genes. Surprisingly, we found these DE genes were enriched in the GO terms and KEGG pathways associated to cancers. These results suggest that U1 snRNA regulates cancer gene expression. Here, we report these results for the first time to facilitate advancements in the further study of U1 snRNA’s functions.

## Results

### U1 snRNA over-expression by transfection

Three U1-transfected samples in the treatment group and three samples in the control group were used to validate the model of U1 snRNA over-expression in PC-12 cells (see methods). After the exogenous U1 snRNA genes expressed for 8 hours in the cells, the qPCR and *in situ* hybridization were conducted for model validation. The data obtained from qPCR assays showed a dose-dependent increase of U1 snRNA in U1-transfected samples, indicating that U1 snRNA had been over-expressed in PC-12 cells (**Figure 1A**). Since transfection with 5 μg vectors resulted in cell death, *in situ* hybridization was conducted to determine the locations of the U1 snRNA transcripts in cells using 4 μg vectors for each sample. Fluorescence images from the control samples indicated U1 snRNA transcripts were located in nuclei under normal conditions, while U1 snRNA transcripts had cytoplasmic accumulation in U1-transfected samples (**Figure 1B**).

**Figure 1.**
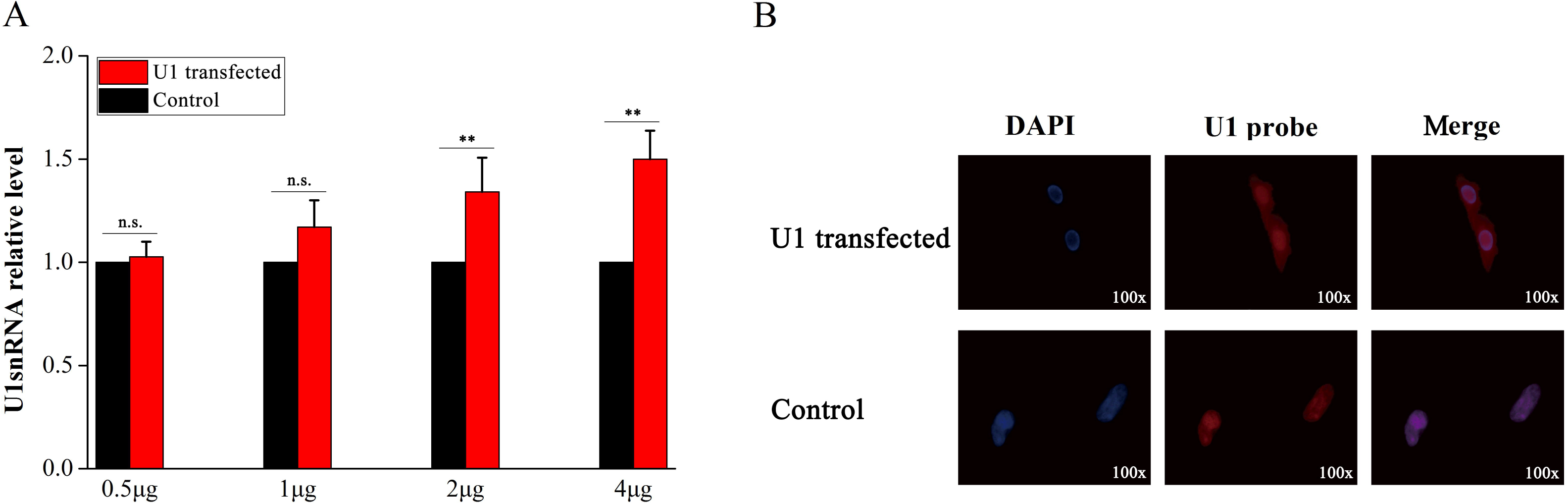
Validation of the U1 snRNA over-expression in PC-12 cells. Three samples in the treatment group were transfected with pSIREN-RetroQ vectors containing the U1 snRNA sequences separately. The other three samples in the control group were transfected with empty pSIREN-RetroQ vectors containing 5 bp polyA. After the exogenous U1 snRNA genes expressed for 8 hours in the cells, the qPCR and *in situ* hybridization were conducted for model validation. **A.** The expression levels of U1 snRNA in six samples were measured by qPCR and the expression levels in three control samples were normalized into 1. **B.** 4μg U1 snRNA vectors or 4μg empty vectors were used for transfection. Cell nuclei (in blue color) were stained with DAPI (in the left column). U1 snRNA were *in situ* hybridized with biotin-labeled LNA probes stained in red color (in the middle column).

### Differentially expressed genes and GO annotation

Three U1-transfected samples using 4 μg vectors and three control samples were used for the microarray experiments. In total, 916 (about 4.42 % of 20,715) DNA probes were identified significantly differentially expressed between the treatment (U1-transfected) group and the control group. These probes represented 595 up-regulated and 321 down-regulated known genes or loci (**Supplementary 1**). These 916 significantly differentially expressed (DE) genes were annotated by Gene Ontology categories (GO terms) in three GO domains (**Supplementary 2**), which were molecular function (MF), cellular component (CC) and biological process (BP). In the MF domain, highly represented categories were cytokine activity (GO:0005125), chemokine activity (GO:0008009), chemokine receptor binding (GO:0042379), and so on. The top three enriched categories in the CC domain were extracellular space (GO:0005615), extracellular region part (GO:0044421), extracellular region (GO:0005576). In the BP domain, they were response to wounding (GO:0009611), regulation of phosphorylation (GO:0042325), regulation of transferase activity (GO:0051338). The enrichment analysis showed tumor necrosis factor receptor superfamily binding (GO: 0032813) and tumor necrosis factor receptor binding (GO: 0005164) in the MF domain were associated to cancers.

### Cancer related pathways and further analysis

The 916 DE genes were also annotated by Kyoto Encyclopedia of Genes and Genomes (KEGG) database (**Supplementary 3**). The enrichment analysis resulted in 12 enriched pathways in total (**Table 1**). Among these 12 pathways, three well-known cancer pathways were p53 signaling pathway (KEGG: rno04115), bladder cancer (KEGG: rno05219) and pathways in cancer (KEGG: rno05200). As for the other nine pathways, MAPK signaling pathway (KEGG: rno04010) [8], cell cycle pathway (KEGG: rno04110) [9], cytokine-cytokine receptor interaction pathway (KEGG: rno04060) [10], NOD-like receptor signaling pathway (KEGG: rno04621) [11], cytosolic DNA-sensing pathway (KEGG: rno04623) [12], RIG-I-like receptor signaling pathway (KEGG: rno04622) [13], Toll-like receptor signaling pathway (KEGG: rno04620) [14], chronic myeloid leukemia pathway (KEGG: rno05220) [15] and Hematopoietic cell lineage pathway (KEGG: rno04640) [16] were also reported to be associated to cancers in the previous studies. To confirm the results from the microarray data analysis, the expression levels of six out of the total 21 DE genes (**Figure 2**) in pathways in cancer (KEGG: rno05200) were measured using the qPCR. The fold changes of six DE genes between the treatment and the control samples showed that Myc, Fos and Nfkb1 had been up-regulated and Ccne1, Fgf2 and Tgfb1 had been down-regulated, which was consistent with the results from the microarray experiments (**Figure 3**). Among the total 73 genes involved in 12 enriched pathways, 56 and 17 were up- and down-regulated, respectively (**Supplementary 3**).

**Table 1.** Enriched pathways induced by the U1 snRNA over-expression

| Pathway | Up-regulated | Down-regulated |
| --- | --- | --- |
| rno04010:MAPK signaling pathway | Pak1, Nf1, Il1r2,<br>Rras2, Dusp8, Nfkb1,<br>Rapgef2, Fos, Traf2,<br>Dusp5, Myc, Ddit3,<br>Tgfb2, Tnf, Gadd45a,<br>Gadd45g, Dusp1,<br>Gadd45b | Pla2g2a, Tgfb1, Cd14,<br>Map4k2, Cdc25b, Ntf4,<br>Pla2g6, Fgf2, Ntrk2 |
| rno04110:Cell cycle | Wee2, Cdkn1a, Smc1b,<br>Mdm2, Cdk6, Cdkn2a,<br>Myc, Tgfb2, Gadd45a,<br>Gadd45g, Gadd45b | Tgfb1, Rbl2, Cdc6,<br>Cdc25b, Ccne1 |
| rno04060:Cytokine-cytokine receptor interaction | Il1r2, Tnfsf15, Il9r,<br>Ccl5, Ifnb1, Cxcl2,<br>Tnf, Il22ra2, Csf1,<br>Cx3cl1, Cxcl10, Ltb,<br>Tnfsf18, Il23r, Il6,<br>Ccl4, Cxcl11 | Tnfsf9, Il23a, Csf3 |
| rno04621:NOD-like receptor signaling pathway | Ripk2, Cxcl1, Nfkb1,<br>Birc3, Ccl5, Tnfaip3,<br>Cxcl2, Tnf, Il6 | Pycard |
| rno04115:p53 signaling pathway | Bid, Cdkn1a, Mdm2,<br>Cdk6, Cdkn2a, Gadd45a,<br>Gadd45g, Bbc3,<br>Gadd45b | Ccne1 |
| rno04623:Cytosolic DNA-sensing pathway | Nfkb1, Ccl5, Ifnb1,<br>Cxcl10, Il6, Ccl4 | Pycard |
| rno04622:RIG-I-like receptor signaling pathway | Ifih1, Nfkb1, Traf2,<br>Ifnb1, Tnf, Dhx58,<br>Isg15, Cxcl10 |  |
| rno04620:Toll-like receptor signaling pathway | Nfkb1, Fos, Tlr5,<br>Ccl5, Ifnb1, Tnf,<br>Cxcl10, Il6, Cxcl11 | Cd14 |
| rno05220:Chronic myeloid leukemia | Cdkn1a, Nfkb1, Mdm2,<br>Runx1, Cdk6, Cdkn2a,<br>Myc, Tgfb2 | Tgfb1 |
| rno04640:Hematopoietic cell lineage | Il1r2, Cd22, Il9r,<br>Tnf, Csf1, Il6 | Cd14, Csf3, Cd37 |
| rno05219:Bladder cancer | Cdkn1a, Cdh1, Mdm2,<br>Rassf1, Cdkn2a, Myc |  |
| rno05200:Pathways in cancer | Bid, Cdkn1a, Cdh1,<br>Nfkb1, Fos, Mdm2,<br>Runx1, Hhip, Traf2,<br>Rassf1, Cdk6, Birc3,<br>Cdkn2a, Myc, Traf1,<br>Tgfb2, Lamc2, Il6 | Tgfb1, Ccne1, Fgf2 |

Pathways used the data from the KEGG database. The gene names used the HUGO gene symbols

**Figure 2.**
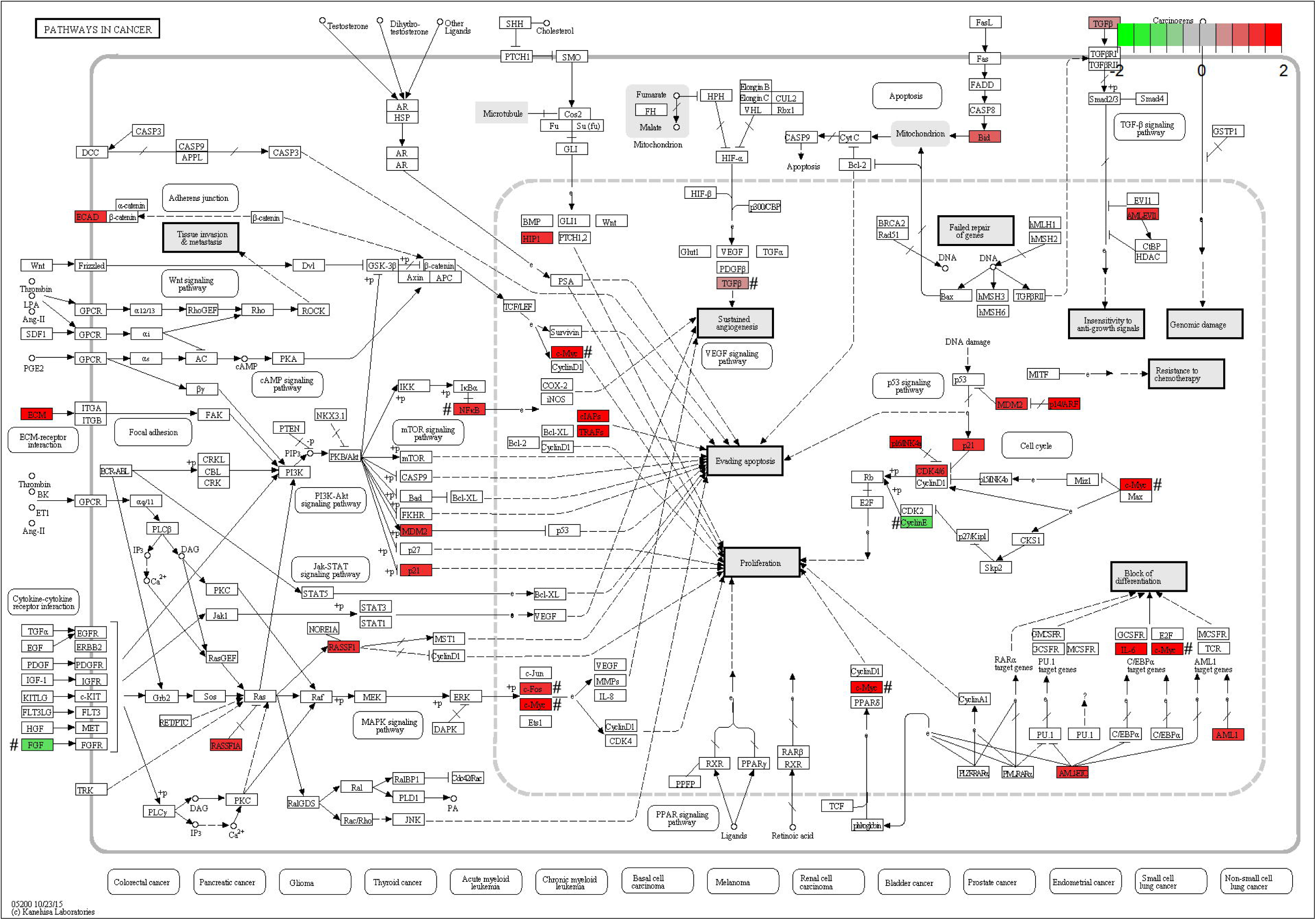
Pathways in cancer. The figure of Pathways in cancer (KEGG: rno05200) was downloaded from the KEGG website. In this map, 21 DE genes included 18 up-regulated genes (Cdk6, Cdkn1a, Lamc2, Il6, Myc, Cdkn2a, Hhip, Traf2, Fos,Mdm2, Rassf1, Runx1, Bid, Traf1, Birc3, Nfkb1, Tgfb2 and Cdh1) in red color and three down-regulated genes (Ccne1, Fgf2 and Tgfb1) in green color. # The expression levels of six genes were confirmed by qPCR.

**Figure 3.**
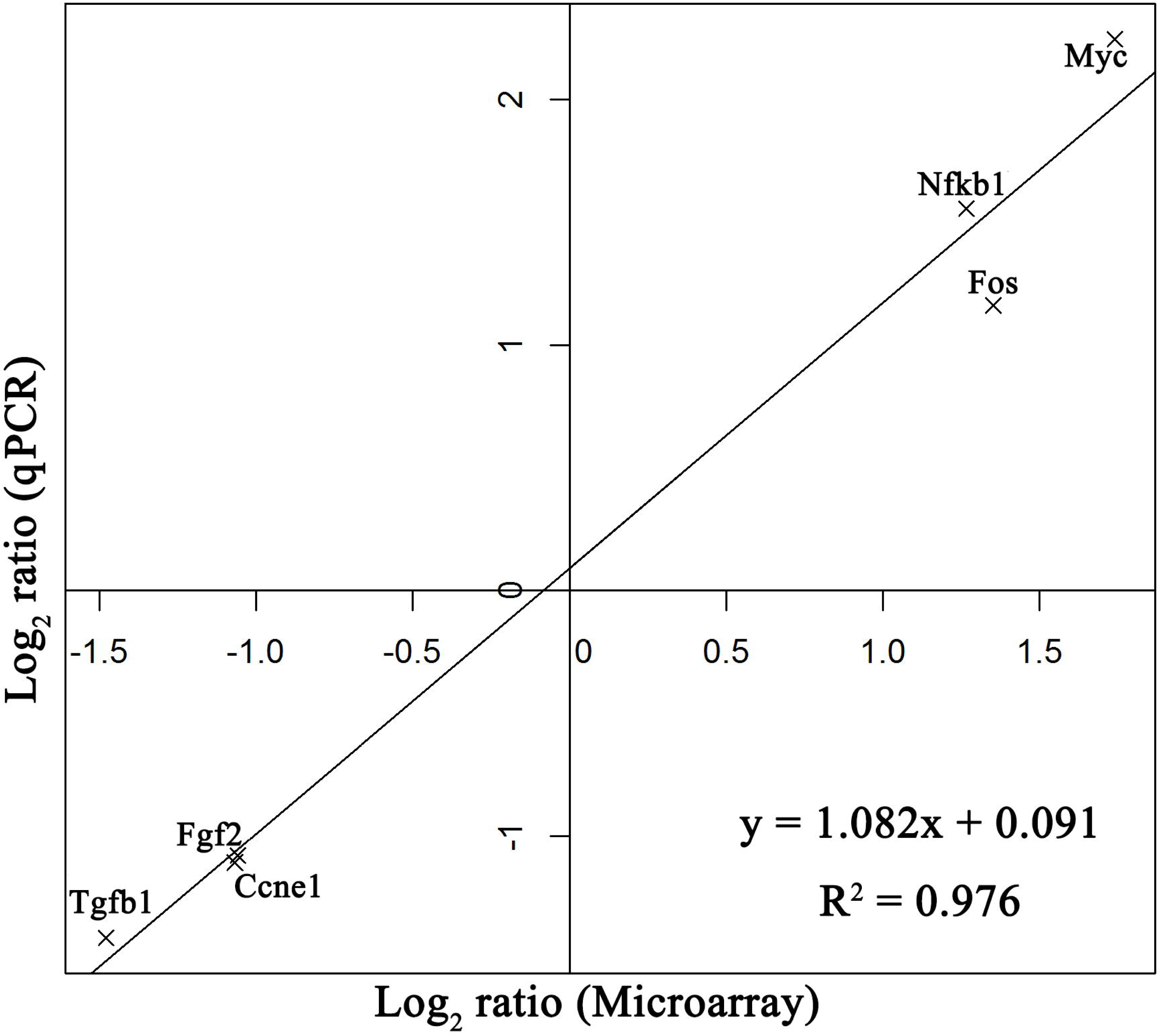
Confirmation of microarray results by qPCR. The expression levels of three up-regulated DE genes (Myc, Fos and Nfkb1) and three down-regulated DE genes (Ccne1, Fgf2 and Tgfb1) were measured using the qPCR. The log2 ratio represents the log2 fold changes of six DE genes between the treatment and the control samples. A linear model was fit using the data of six genes.

The results in this study suggest U1 snRNA regulates the gene expression, which can be explained by the theory of premature termination by cleavage and polyadenylation (PCPA) [7]. The PCPA process yields shorter mRNAs using more proximal alternative polyadenylation (APA) sites, which has already been proved in activated immune, neuronal, and cancer cells [7]. The U1 snRNA over-expression in PC-12 neuronal cells resulted in more longer mature mRNAs with more RNA-binding or protein-binding domains for regulation. From our observations, we found another mechanism to explain U1 snRNA regulates the gene expression. The over-abundant exogenous U1 snRNA transcripts are transported to cytoplasm (**Figure 1B**) but some of them cannot assembled into U1 snRNPs. The local double-stranded RNAs (dsRNAs) on these unassembled U1 snRNAs can trigger the RNA interference (RNAi) in cytoplasm and produce small interfering RNAs (siRNAs) [17]. In brief, U1 snRNA could regulate the gene expression through RNAi.

## Materials and methods

### Transfection of U1 snRNA into PC-12 cells

PC-12 cells, derived from a pheochromocytoma of the rat adrenal medulla were cultured in RPMI-1640 medium containing 10% fetal bovine serum in our lab. The medium was changed every two days and the cells were passaged every four days. To validate the U1 snRNA over-expression by transfection, PC-12 cells were sampled for six times (about 2×10^6^ cells per sample), then trypsinized, washed once with PBS and resuspended in 100 μL Nucleofector Solution (Lonza-Amaxa, German). Three samples (treatment group) were transfected with 0.5, 1, 2, 4 and 5 μg pSIREN-RetroQ vectors [7] (Clontech, USA) containing the 168 bp U1 snRNA sequences (Genbank: V01266.1) separately. The other three samples (control group) were transfected with empty pSIREN-RetroQ vectors containing 5 bp polyA as control. The transfection was performed by electroporation using a Lonza-Amaxa Nucleofector Pulser with the program setting U-029. Six samples were cultured in 2.5 mL RPMI-1640 medium for 8 h. Total RNA was isolated from six samples using the RNAiso Plus Reagent (TaKaRa, Japan) and then processed separately. The cDNA of U1 snRNA was synthesized by Mir-X™ miRNA First-Strand Synthesis Kit (Clontech, USA). The cDNA product was amplified by qPCR (Eppendorf, German) using U6 snRNA as internal control (**Supplementary 4**). The qPCR reaction mixture was incubated at 95°C for 30 s, followed by 40 PCR cycles (5 s at 95°C and 20 s at 62°C for each cycle).

The U1 snRNA transcripts in six samples were *in situ* hybridized by 22 bp biotin-labeled LNA probes 5′-GTATCTCCCCTGCCAGGTAAGT-3′ (Exiqon, Denmark) following the procedure below. For each sample, the hybridization buffer (50% v/v formamide, 2X SSC, 50 mM sodium phosphate with pH 7 and 10% dextran sulfate) were mixed with 10 nM probes, and then added into the cells at 55°C for 12 h in a humidified chamber. After hybridization, cells were washed in 2X SSC, then incubated in 0.1% Triton at 4°C for half an hour, and washed in 2X SSC. A fluorescent Alexa Fluor 594 streptavidin conjugate (Yeasen, China) was added into the cells to stain biotin-labeled LNA probes to the red color. Cells were washed in 4X SSC with 0.1% Triton at 4°C, and then washed in 2X SSC, 1X SSC and PBS subsequently at room temperature. Cell nuclei in six samples were stained by DAPI to the blue color and examined under a fluorescence microscope (Olympus, Japan).

### Microarray experiments

The second generation of PC-12 cells were passaged to conduct microarray experiments. These PC-12 cells were sampled for six times (about 2×10^6^ cells per sample). Three samples in the treatment group and the other three samples in the control group were processed following the same procedure described above. Total RNA was isolated from six samples using RNAiso Plus Reagent (TaKaRa, Japan) and then processed separately. The cDNA synthesis and antisense RNA (aRNA) amplification were conducted using Amino Allyl MessageAmp II aRNA Amplification Kit (Ambion, USA). The modified nucleotide, 5-(3-aminoallyl)-UTP (aaUTP) was incorporated into the amplified aRNA during the *in vitro* transcription with T7 RNA to form amino allyl-aRNA (aa-aRNA). The aa-aRNA was readily labeled with amine-reactive N-hydroxysuccinimide (NHS) ester of fluorescent dyes (Cy5-NHS). Finally, six samples were hybridized to six chips of Rat OneArray™ Plus (Phalanx Biotech, China) separately. The Rat OneArray™ Plus chip contains 20,715 DNA probes with the length of 60 bp.

### Confirmation of microarray results by qPCR

The third generation of PC-12 cells were passaged to confirm the changes of significantly differentially expressed (DE) genes selected using the microarray data. These PC-12 cells were sampled for six times (about 2×10^6^ cells per sample). Three samples in the treatment group and the other three samples in the control group were processed for U1 snRNA transfection, RNA extraction and cDNA amplification, following the same procedure described in the first paragraph of this section. The qPCR reaction mixture was incubated at 95°C for 30 s, followed by 40 PCR cycles (5 s at 95°C and 30 s at 60°C for each cycle).

### Data analysis

Six chips were scanned into images in TIF format using the Agilent G2505C Microarray Scanner System (Agilent Technology, USA). Images were read into gene expression data (numerical values) in GPR format using the software GenePix Pro v4.1.1.44 (Axon Instruments, CA). These gene expression data were normalized and further analyzed to determine the significantly differentially expressed (DE) genes between the treatment group and the control group using the software Rosetta Resolver System v7.2 (Rosetta Biosoftware, USA). All the DE gene satisfied both the P-value < 0.05 and the absolute log2 fold-change >= 1. DNA probes were annotated by the software Annotation ROA2 r1.0 (Phalanx Biotech, China) with two databases NCBI RefSeq r65 and Ensembl r76 cDNA sequences Rnor_5.0 annotations. Statistical computing and graphics were implemented using R v2.15.3[18]. The annotation and functional enrichment analyses of GO and KEGG pathways were implemented using WebGestalt (http://www.webgestalt.org/).

## Acknowledgments

The data analysis in this study was supported by National Scientific Data Sharing Platform for Population and Health Translational Cancer Medicine Specials.

## Authors' contributions

TZ, SG and ZC conceived and supervised this project. ZC and YCS performed the experiments. SG, YS, JS and ZW analyzed the data. SG and ZC drafted the manuscript. SG modified and polished the manuscript. XN prepared the figures, tables and supplementary materials. XN managed the manuscript submission.

